# The genetics of male pheromone preference difference between *Drosophila melanogaster* and *D. simulans*

**DOI:** 10.1101/793315

**Authors:** Michael P. Shahandeh, Alison Pischedda, Jason M. Rodriguez, Thomas L. Turner

## Abstract

Species of flies in the genus *Drosophila* differ dramatically in their preferences for mates, but little is known about the genetic or neurological underpinnings of this evolution. Recent advances have been made to our understanding of one case: pheromone preference evolution between the species *D. melanogaster* and *D. simulans*. Males of both species are very sensitive to the pheromone 7,11-HD that is present only on the cuticle of female *D. melanogaster*. In one species this cue activates courtship, and in the other it represses it. This change in valence was recently shown to result from the modification of central processing neurons, rather than changes in peripherally expressed receptors, but nothing is known about the genetic changes that are responsible. In the current study, we show that a 1.35 Mb locus on the X chromosome has a major effect on male 7,11-HD preference. Unfortunately, when this locus is divided, the effect is largely lost. We instead attempt to filter the 159 genes within this region using our newfound understanding of the neuronal underpinnings of this phenotype to identify and test candidate genes. We present the results of these tests, and discuss the difficulty of identifying the genetic architecture of behavioral traits and the potential of connecting these genetic changes to the neuronal modifications that elicit different behaviors.

## Introduction

Understanding the proximate mechanisms of phenotypic divergence has long been a goal of evolutionary biologists (Stern et al., 2009; Stern & Orgogozo, 2008). Advances in genome sequencing have led to a recent boom in genotype-phenotype association studies, the large majority being for morphological traits (Orgogozo & Martin, 2013). These findings have resulted in a better understanding of the molecular underpinnings of morphological evolution, allowing us to observe general patterns about the types of genetic changes commonly associated with phenotypic change, the number of loci involved, and the general size of their effects (Kittelmann et al., 2017; Martin & Orgogozo, 2013; Rebeiz & Williams, 2017). Comparatively, we have fewer studies of the proximate mechanisms underlying behavioral divergence with which to draw broad conclusions. This is unfortunate, because behaviors are important phenotypes, particularly with respect to speciation and biodiversity. For example, differences in host, habitat, and mating behaviors can form strong reproductive barriers between species (Coyne & Orr, 1997, 2004).

Behaviors holistically involve the detection of stimuli via the peripheral nervous system, sensory integration via central nervous system processing, and the coordinated production of a behavioral output. It is therefore surprising that a large proportion of genes known to cause behavioral divergence between species affect sensory perception at the periphery, rather than the other molecular determinants of behavior (Auer et al., 2019; Cande et al., 2013; Leary et al., 2012; McBride et al., 2014). This could indicate that changes at the periphery, mainly in membrane-bound stimulus-detecting receptors, are favored targets of selection because they can have drastic effects on specific phenotypes while minimizing pleiotropic effects (McBride, 2007). It is, however, difficult to generalize from so few examples. Moreover, it is quite possible that this pattern is due mainly to ascertainment bias (Rockman, 2012). Indeed, many case studies that have successfully mapped causal genes explaining behavioral divergence have used a candidate-gene approach targeting sensory receptors, so it is plausible that other types of changes are more common but have been overlooked. For example, recent studies that take whole-genome approaches have identified important variants in genes affecting behavior that act at the synapse (Kocher et al., 2018), or as hormonal neuromodulators (Bendesky et al., 2017). To make generalized predictions about the types of changes underlying behavioral divergence, we need more studies of the genetic basis of behavior that take unbiased approaches, preferably in systems with multiple comparable cases of evolution in similar behaviors (i.e. “metamodel systems” sensu Kopp, 2009). With an active research community, many genetic tools, and an easily manipulated life-history, the *Drosophila* species group provides an excellent opportunity to make further progress, particularly because closely related species differ dramatically in many behaviors.

Courtship preference is one behavior that varies dramatically among *Drosophila* species. In *Drosophila*, males and females express a blend of cuticular hydrocarbons (CHCs), many of which act as gustatory pheromones (Pardy et al., 2018). These CHCs are variable between species (Jallon & David, 1987), and can act as sex pheromones (Ferveur & Sureau, 1996) and species identification signals (Billeter et al., 2009). For example, *D. melanogaster* females predominantly express 7,11-heptacosadiene (7,11-HD), while females in the closely-related species, *D. simulans*, primarily express 7-tricosene (7T). The male responses to these pheromones differ dramatically between species: *D. melanogaster* males willingly court conspecific females, but the presence of 7,11-HD on the *D. melanogaster* female cuticle suppresses courtship by *D. simulans* males (Billeter et al., 2009; Clowney et al., 2015; Seeholzer et al., 2018). Male pheromone preference, therefore, constitutes an early barrier to interspecific mating (Shahandeh et al., 2018).

Both *D. melanogaster* and *D. simulans* males detect 7,11-HD via gustatory receptor neurons in the forelegs that express the ion channel *ppk23* (Lu et al., 2012, 2014; Thistle et al., 2012; Toda et al., 2012). From there, the signal is propagated through two clusters of neurons (vAB3 and maL neurons) that simultaneously excite and inhibit the P1 central courtship neurons (Clowney et al., 2015). These P1 central courtship neurons act as command neurons, essentially like an on/off switch for male courtship (Auer & Benton, 2016). A recent study found that the evolution of the interactions among these neurons in the central nervous system causes the difference in 7,11-HD preference, rather than the evolution of the peripheral nervous system. (Seeholzer et al., 2018). In *D. melanogaster*, 7,11-HD promotes courtship because the excitation of P1 neurons is greater than the inhibition; in *D. simulans*, the opposite seems to be the case (Seeholzer et al., 2018). However, the molecular changes underlying these differences in neuronal interactions remain unknown. This detailed, if still incomplete, understanding of the cellular basis of behavioral evolution presents an excellent opportunity to map the causal genes and link evolution in behavior at the genetic, cellular, and organismal levels.

Here we present the results of a genotype-phenotype association study where we make considerable progress toward identifying the loci underlying shifts in 7,11-HD preference behavior between *D. simulans* and *D. melanogaster*. First, we confirm a previous result demonstrating that a portion of the male preference phenotype maps to the X chromosome (Kawanishi & Watanabe, 1981). We then show that the preference for 7,11-HD can be recovered in hybrids with a single 1.35 Mb region of the *D. melanogaster* X chromosome. We additionally present the results of two attempts to map this region to a causal gene; both a fine-mapping and candidate gene approach were unsuccessful. Nonetheless, our findings have identified a fraction of the *D. melanogaster* genome containing loci for further functional investigation.

## Methods

### Fly stocks and maintenance

We maintained all fly strains in 25 mm diameter vials on standard cornmeal/molasses/yeast medium at 25°C under a 12 h:12 h light/dark cycle. Under these conditions, we established non-overlapping two-week lifecycles. Every 14 days, we transferred all of the emerged male and female adult flies into vials containing fresh food, where they were allowed to oviposit for 1–3 days before being discarded. To test for species differences in male preference, we used two *D. simulans* strains: simC167.4 (obtained from the UC San Diego *Drosophila* Stock Center; Stock #: 14021-0251.99) and *Lhr* (Brideau et al., 2006; Watanabe, 1979). We used four *D. melanogaster* strains: Canton-S, DGRP-380 (MacKay et al., 2012), C(1)DX-LH_M_ and LH_M_ (Rice et al., 2005). To screen portions of the X chromosome, we created duplication hybrids (see below) using a total of 22 Dp(1;Y) strains listed in Table 1 (Cook et al., 2010).

**Table 1.**
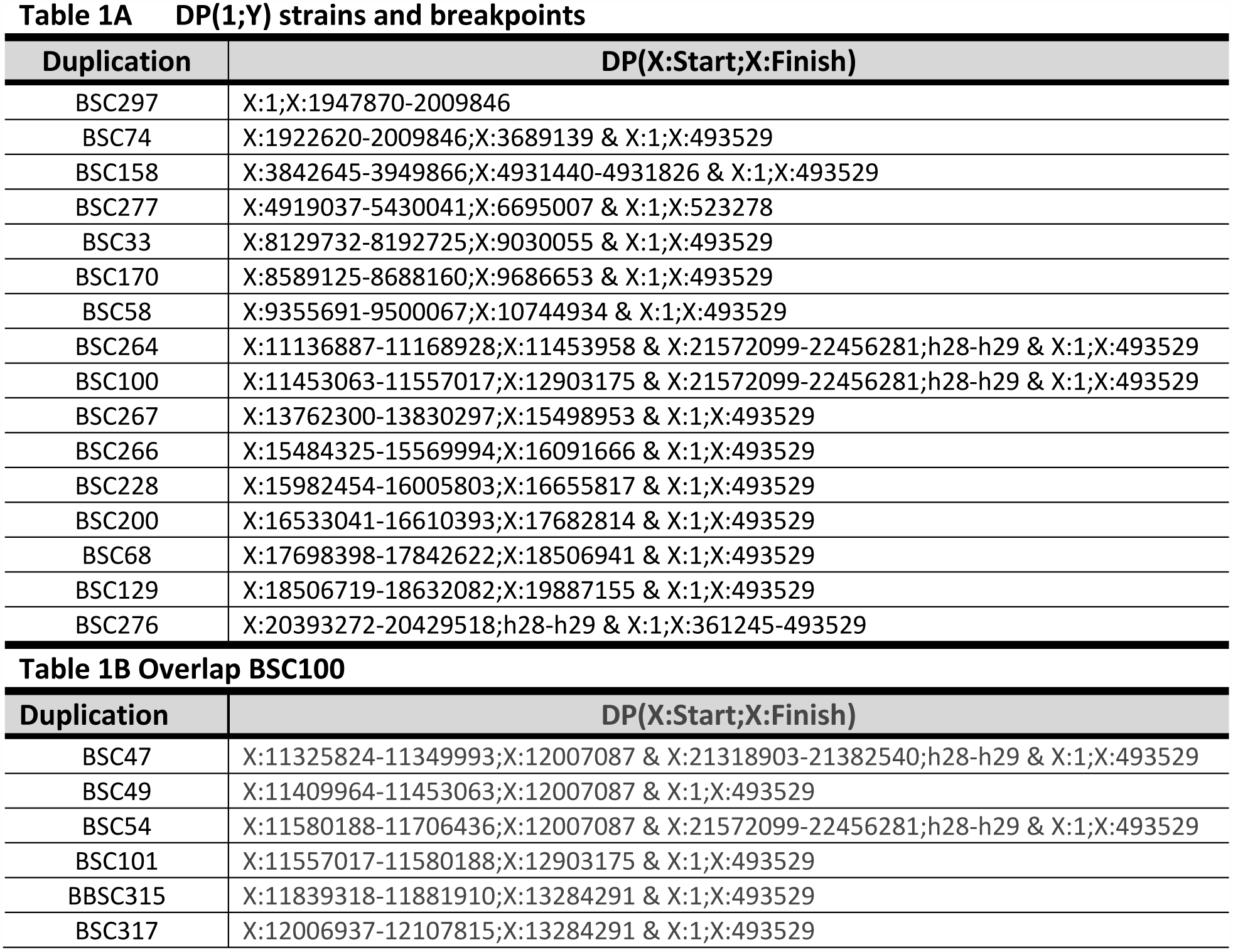
DP(1;Y) *D. melanogaster* stocks. **1A.** The 16 Y-linked X duplication strains used to create duplication hybrids and the reported breakpoints of their Y-linked X duplication segment (if multiple segments, base pairs are indicated for each). Ranges are shown for breakpoints where precise estimation is not available. **1B.** The 6 Y-linked X duplication strains used to create duplication hybrids overlapping the region covered by BSC100 and the reported breakpoints of their Y-linked X duplication segment. Note, all translocations contain a basal segment of the X chromosome (X:1;X:493529), and many also contain a region from the end of the X chromosome (ex: X:21572099-22456281). Coordinates of translocated segments are from *D. melanogaster* genome release 6 (Dos Santos et al., 2015).

### Hybrid crosses

To confirm the role of the X chromosome in male mate discrimination, we needed to make F1 hybrid males by crossing *D. melanogaster* females to *D. simulans* males (hybrid offspring would then have a *D. melanogaster* X) and by the reciprocal cross (hybrids would have the same autosomal genotype, but the *D. simulans* X). *D. melanogaster*/*D. simulans* hybrid males normally die during development, while hybrid females are infertile (Watanabe, 1979). The *Lhr* strain of *D. simulans* rescues male viability, allowing us to collect living male and female hybrid offspring (Figure 1A). However, crossing females from this *D. simulans* hybrid male rescue strain to *D. melanogaster* males never yielded offspring, probably because of very strong pre-mating isolation. Instead, we used genetic tools (see below) to make these same genotypes while only crossing *D. melanogaster* females to *D. simulans* males. In this crossing direction, we found hybrids could be made using the following steps. First, we collected 20 *D. simulans* males as virgins and aged them for 7-12 days. We collected 10 *D. melanogaster* females as very young virgins, just 2-4 hours after eclosion. We immediately combined 10 very young *D. melanogaster* virgins with 20 aged *D. simulans* males in a vial with food media. We pushed a foam plug into the vial, leaving only 1-2 cm of space above the food surface. We held these hybrid cross vials in this manner for 2-3 days before transferring the flies to a new vial with fresh food to oviposit.

**Figure 1.**
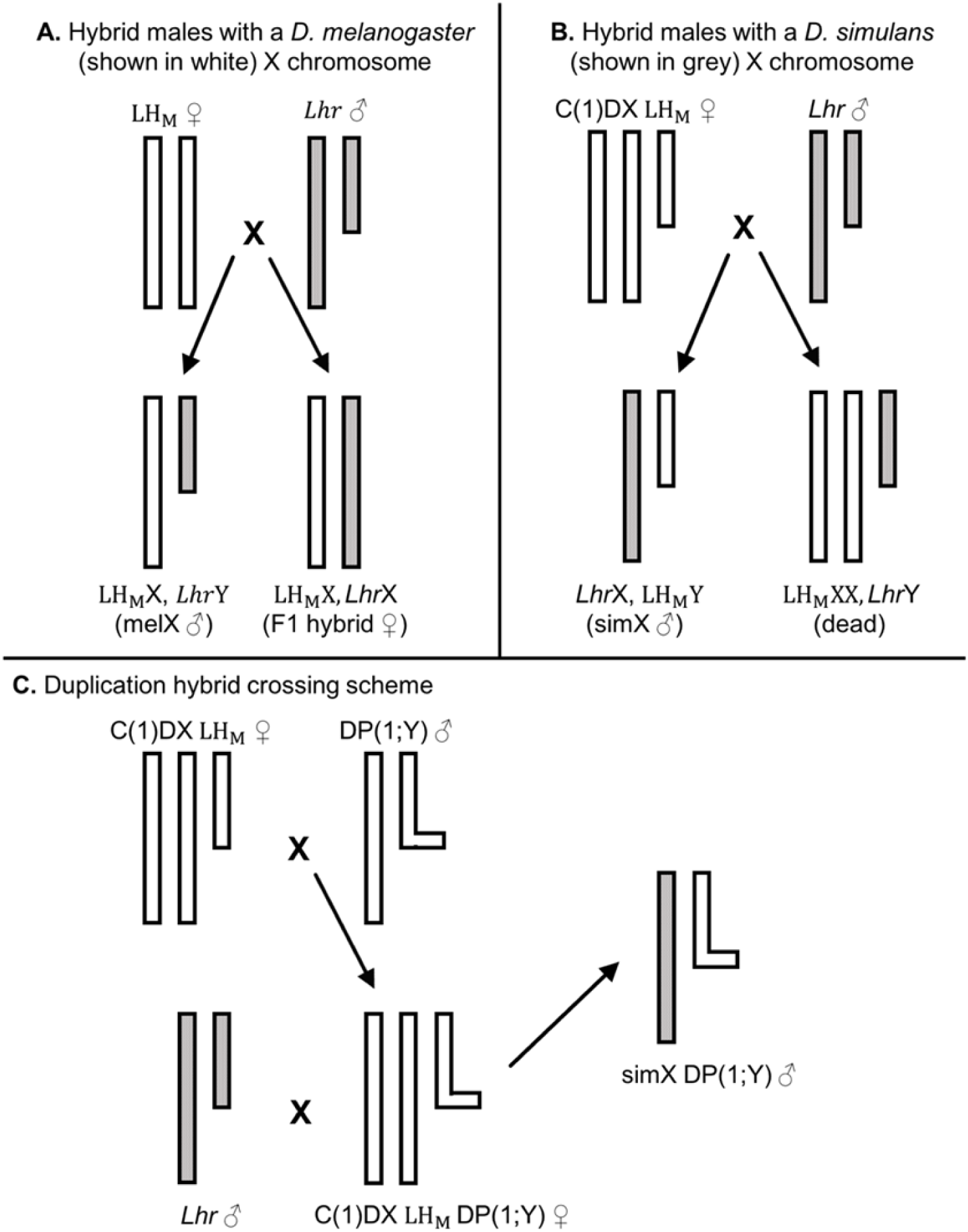
Hybrid crossing schemes. Each diagram shows only the sex chromosomes. Y chromosomes are depicted as shorter, while Y chromosomes with X duplications are depicted as an “L”. *D. melanogaster* (LH_M_) chromosomes are shown in white, and *D. simulans* (*Lhr*) chromosomes are shown in grey. **A.** To create melX hybrids, we crossed LH_M_ females to *Lhr* males, resulting in hybrid males with the *D. melanogaster* X chromosome and F1 hybrid females. **B.** To create simX hybrids, we crossed C(1)DX-LH_M_ females to *Lhr* males, resulting in hybrid males with the *D. simulans* X chromosome. Females of this cross are inviable. **C.** To create duplication hybrids, we first crossed males from DP(1;Y) strains to C(1)DXLH_M_ females. We took the female offspring of this first cross and crossed them to *Lhr* males. The male offspring of this second cross inherit the *D. simulans* X chromosome in addition to a segment of the *D. melanogaster* X chromosome (depicted by the small horizontal bar) attached to the *D. melanogaster* Y chromosome.

To create male hybrids with the *D. melanogaster* X chromosome (melX), we set up the above crosses using the LH_M_ strain of *D. melanogaster* and the *Lhr* strain of *D. simulans*. To create male hybrids with the X chromosome of *D. simulans* (simX), we set up the same cross using the C(1)DX-LH_M_ strain of *D. melanogaster* and the *Lhr* strain of *D. simulans*. Females of the C(1)DX-LH_M_ strain have a compound X chromosome and a Y chromosome in an LH_M_ autosomal background. The male offspring of this cross inherit their X chromosome from the father, while their Y chromosome and cytoplasm are inherited from the mother (Figure 1B). Thus, simX and melX hybrids only differ in their sex chromosomes. By directly comparing them we can isolate the effect of the sex chromosomes (primarily the X chromosome) on male courtship preference behavior.

### Male courtship assays

For all courtship assays, we aged virgin males and females in single-sex vials for 4 days at 25°C in densities of 10 and 20, respectively. We gently aspirated a single experimental male into a 25 mm diameter vial with standard cornmeal/molasses/yeast medium 24 hours before observation. On the morning of observation, we aspirated a single female into the vial and pushed a foam plug down into the vial, leaving a space of 1-3 cm above the food surface. This limited space ensures that flies interact within the observation time period. We observed flies for 30 minutes, collecting minute-by-minute courtship data by manually scoring each pair for three easily observed stages of courtship: singing (single wing extensions and vibration), attempted copulation, and successful copulation. We scored pairs that exhibited multiple stages within a single minute once within that minute. We conducted 2 observations per day at room temperature between the hours of 9 and 11 AM (0-2 hours after fly incubator lights turn on). All observers were blind to both male and female genotype (see below).

We tested the male courtship behavior of the following strains: *D. melanogaster* (LH_M_, Canton-S, RNAi strains), *D. simulans* (*Lhr* and simC167.4), and various melX and simX F1 hybrids. We measured male courtship towards three types of females, *D. melanogaster, D. simulans*, and F1 hybrid females, using no-choice (single female) assays. All *D. simulans* female courtship objects were of the simC167.4 strain. Female courtship objects for *D. melanogaster* were of the DGRP-380 strain when testing males of the LH_M_, *Lhr*, melX, simX, and duplication hybrid genotypes (i.e. when testing for an effect of the X chromosome), but we used females from the Canton-S strain with Canton-S males, RNAi males and controls, and gene aberration melX hybrids (see below), because these additional data were collected during a separate follow-up experiment. We made F1 hybrid females using the methods described above (i.e. from a cross between LH_M_ and *Lhr*; Figure 1A).

### Duplication hybrid crosses

Mapping the genes responsible for this evolved behavior is very challenging for several reasons. QTL analysis is not possible using these species, as hybrid males are inviable and hybrid females are infertile. One way around this problem is to use large engineered deletions to make loci hemizygous rather than heterozygous in F1 hybrids, exposing recessive or additive alleles from the *D. simulans* parent (Laturney & Moehring, 2012; Moehring & Mackay, 2004; Pardy et al., 2018; Ryder et al., 2004). However, a deletion screen is not possible for an X-linked behavior present only in males because the X is already hemizygous, so large deletions are lethal. Instead, we used a set of transgenic strains (Cook et al., 2010) to create duplication hybrid males – hybrids with a complete *D. simulans* X chromosome and an additional segment of the *D. melanogaster* X chromosome translocated to *D. melanogaster* Y chromosome. (Figure 1C, Table 1A). This method has been used to map hybrid incompatibility loci (Cattani & Presgraves, 2012), but to our knowledge has not been used to map differences in morphology or behavior. The primary caveat to this method is the inability to detect *D. melanogaster* loci that are recessive to their *D. simulans* counterparts. Because F1 hybrid males are hemizygous, it is impossible to assess dominance a priori. Despite this limitation, the primary advantage to this method is our ability to assay a large portion of the X chromosome (80%) using just 16 DP(1;Y) hybrid strains. To create these hybrids, we crossed *D. melanogaster* males from a DP(1;Y) strain to *D. melanogaster* females from the C(1)DX-LH_M_ strain (Figure 1C). We then took the resulting female offspring, which carry a *D. melanogaster* compound X chromosome and a *D. melanogaster* Y chromosome that has a translocated segment of the X chromosome, and crossed them to *D. simulans Lhr* males using the hybrid cross methods described above. The DP(1;Y) Y chromosome is marked with the dominant visible *Bar* mutation, so inheritance of this chromosome is easy to track. After assaying our original 16 duplication hybrid strains, we tested additional strains with duplications that partially overlap a region we identified in the initial screen in an attempt to fine-map loci within DP(1;Y) segments of interest (Table 1B).

### Perfuming D. simulans females with 7,***11-HD***

Adapting the methods of Thistle et al. (2012), we perfumed *D. simulans* females with synthetic 7,11-HD to test for a role in courtship behavior for a duplication of interest (BSC100). To perfume females, we placed 20 simC167.4 females into an empty 25 mm diameter vial. We had previously added either 40 μL of ethanol (sham treatment), or 200 or 400 μg of 7,11-HD dissolved in 40 μL ethanol (perfume treatment) to the vial, and allowed the liquid to evaporate. We then vortexed the vials on the highest setting for three 20-second intervals separated by 20 seconds of recovery. We allowed females to recover from vortex mixing for 30 minutes before loading them into courtship vials. We conducted courtship assays as described above. We detected no difference in courtship between our 200 and 400 μg perfuming treatments (χ^2^= 0.083, df = 1, p = 0.773), so we combined them into a single perfume treatment in our final analysis.

### Selecting and testing candidate genes for validation

The BSC100 duplication region that has a significant effect on male courtship behavior in a hybrid background (see results) contains 159 genes (Table S1). In order to select appropriate candidate genes to test for a role in divergent male courtship behaviors, we used modENCODE expression data (Graveley et al., 2011) to identify genes expressed in the *D. melanogaster* central nervous system, a justification set by the findings of Seehozer et al. (2018). We further filtered this list of genes, obtained from FlyBase (Dos Santos et al., 2015), for biological function in nervous system development/function, or transcription factor activity to exclude any genes not specific to the nervous system (i.e. cell maintenance loci). Finally, we selected any of the 25 remaining genes that have known *Fruitless* binding sites or interactions, as *Fruitless* is responsible for the male specific wiring of the central nervous system (Goto et al., 2011; Manoli et al., 2005; Vernes, 2014). This left us with a list of 6 candidate genes. For 4 of the 6 genes, we were able to procure a non-lethal aberration (Table 2A). We crossed females of these strains to *D. simulans Lhr* males to create melX hybrids with individual gene knockouts. Two of these knockouts (*Smr* and *pot*) are held over a balancer chromosome in females, and thus produce two types of melX hybrid males: those carrying a balancer (intact) X chromosome, and those carrying a defective X-linked allele. Unfortunately, *pot* defective hybrid males were not viable, and thus could not be observed (Table S3). For *Smr*, we compared both balancer hybrids and *Smr*^-^ hybrids paired with *D. melanogaster* and *D. simulans* females in the courtship assays described above. For the remaining, unbalanced strains (*Ten-a* and *Pde9*), we compared courtship of knockout hybrid males toward *D. simulans* and *D. melanogaster* females. *Smr* and *Pde9* hybrids presented an extra challenge, as males have white eyes (Table 2A), and thus, difficulty tracking a female courtship target. To remedy this challenge, we observed these males as above, but in a smaller reduced arena to increase interaction between visually impaired males and females. To modify the courtship arena for these males, we inserted a single piece of plastic vertically into the media, dividing the vial in half, before pushing a foam plug down into the vial, resting 1-2 cm from the food surface.

**Table 2.**
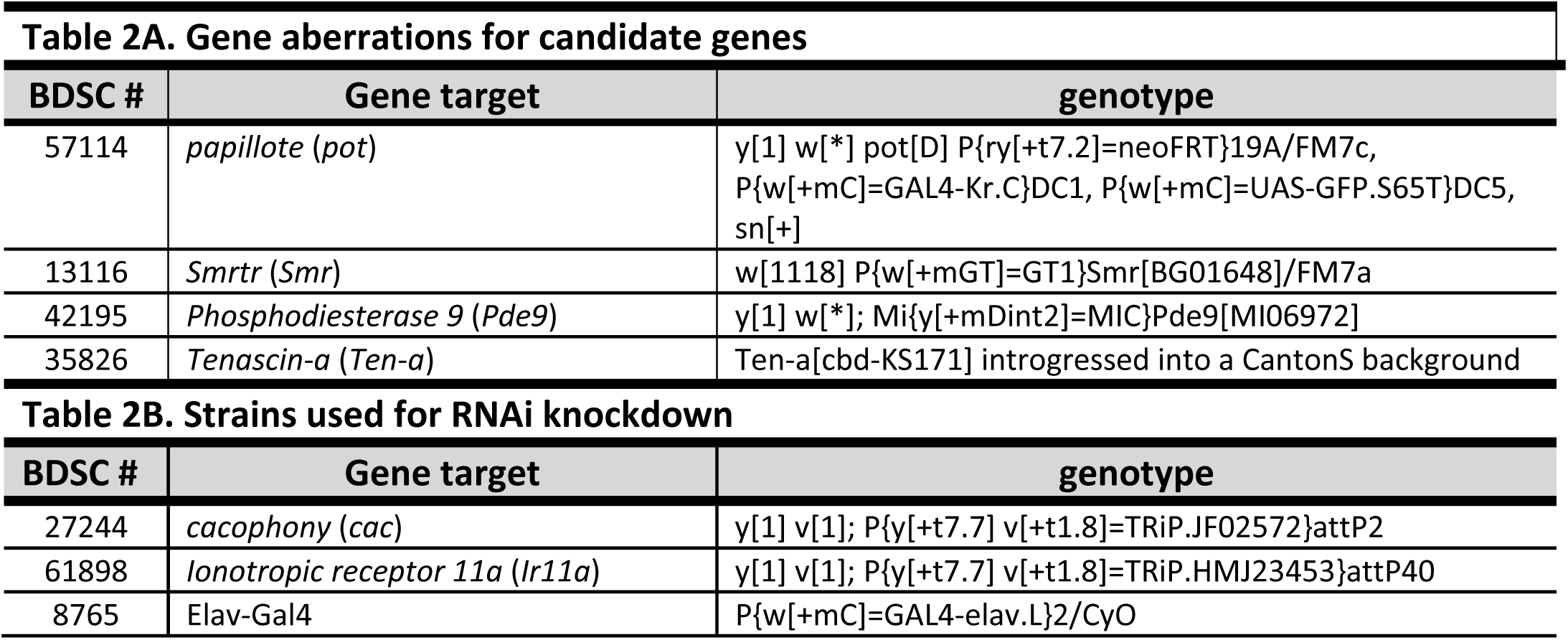
**A.** A list of the available Bloomington *Drosophila* Stock Center gene aberration strains used to test candidate genes. **B.** A list of UAS-hairpin RNAi and pan-neuronal Gal4 drivers used to knockdown expression of genes when aberrations were available, or when aberrations were male lethal.

Because the two remaining genes were either lethal in males (*cacophony*), or had no aberration available at all (*Ir11a*), we knocked down expression in *D. melanogaster* flies using RNAi under the control of the Gal4/UAS system (Perkins et al., 2015). For RNAi knock-down strains, we compared progeny of RNAi lines crossed with the pan-neuronal elav-Gal4 stock (Table 2B). The elav-Gal4 insertion is maintained in heterozygotes over a balancer chromosome. Thus, this cross yields RNAi males, which express hairpin RNAi for the target gene in all of their neurons, and control males, which encode for, but do not express the hairpin RNAi because they lack Gal4 expression in neurons. Control males express a dominant visible marker contained on the balancer chromosome they inherit instead of the Gal4 insertion. In this way, for males paired with both *D. melanogaster* and *D. simulans* females, we compare the courtship of males where expression of the gene of interest is reduced in neurons, to males of the same background without reduced expression in the nervous system. While knocking out (or down) expression of these 6 candidate genes allows us to compare the specific effects of loss of function of *D. melanogaster* alleles to functioning alleles in their respective backgrounds (hybrid for knock-outs, *D. melanogaster* for RNAi knockdown), it is not immediately clear what phenotype to expect during these tests. This presents another challenge in interpreting these results (see Discussion).

### Data analysis

From the minute-by-minute courtship data, for each male-female combination (see below), we collected binomial data (court/did not court) to determine the courtship frequency (CF) of male genotypes. We only considered males that spent 10% or more of the total assay time (i.e., >3 min) in one of the three courtship stages as successfully displaying courtship. For male genotypes, unless otherwise stated, we used Fisher’s exact test to compare the proportion of males that courted a given female type followed by posthoc analysis with sequential Bonferroni tests (Holm, 1979).

For each male that displayed courtship towards a female target, we also calculated the total percent of assay time (30 minutes) that a male spent courting as a proxy for male courtship effort (CE). For males that mated with females, we calculated CE as the percent of time a male spent courting from the start of the assay until the time of copulation. Unlike with CF, we used no minimum threshold for CE. CE is representative of male investment in any given female, another indication of male choice (Edward & Chapman, 2011). Because the courtship effort distributions are highly skewed, we report the median values for comparison across strains. For each male genotype, we compared courtship effort between female genotypes using the Mann-Whitney U test followed by posthoc analysis with sequential Bonferroni tests (Holm, 1979).

## Results

### A significant conspecific courtship preference between D. melanogaster and D. simulans

For both *D. melanogaster* strains that we assayed, we detected significantly higher courtship frequencies when males were paired with females of their own species (Figure 2A, Table S2). Both LH_M_ and Canton-S courted *D. melanogaster* females significantly more frequently than *D. simulans* females (LH_M_: p = 0.0003; Canton-S: p = 0.0032). Likewise, both *D. simulans* strains displayed higher courtship frequencies with *D. simulans* females than *D. melanogaster* females (*Lhr:* p = 5.34E-09; simC167.4: p = 5.47E-12). There were no differences in the courtship frequencies among strains of the same species (p = 1 for both *D. melanogaster* and *D. simulans*). Unsurprisingly, when we calculated a consensus p-value (Rice, 1990), which tests the combined effect of independent tests of the same hypothesis, for the two *D. melanogaster* strains and the two *D. simulans* strains, conspecific courtship preferences remained highly significant (p = 1.61E-5 and p < 1.00E-25, respectively), suggesting this is indeed a species-level, rather than strain-specific difference.

**Figure 2.**
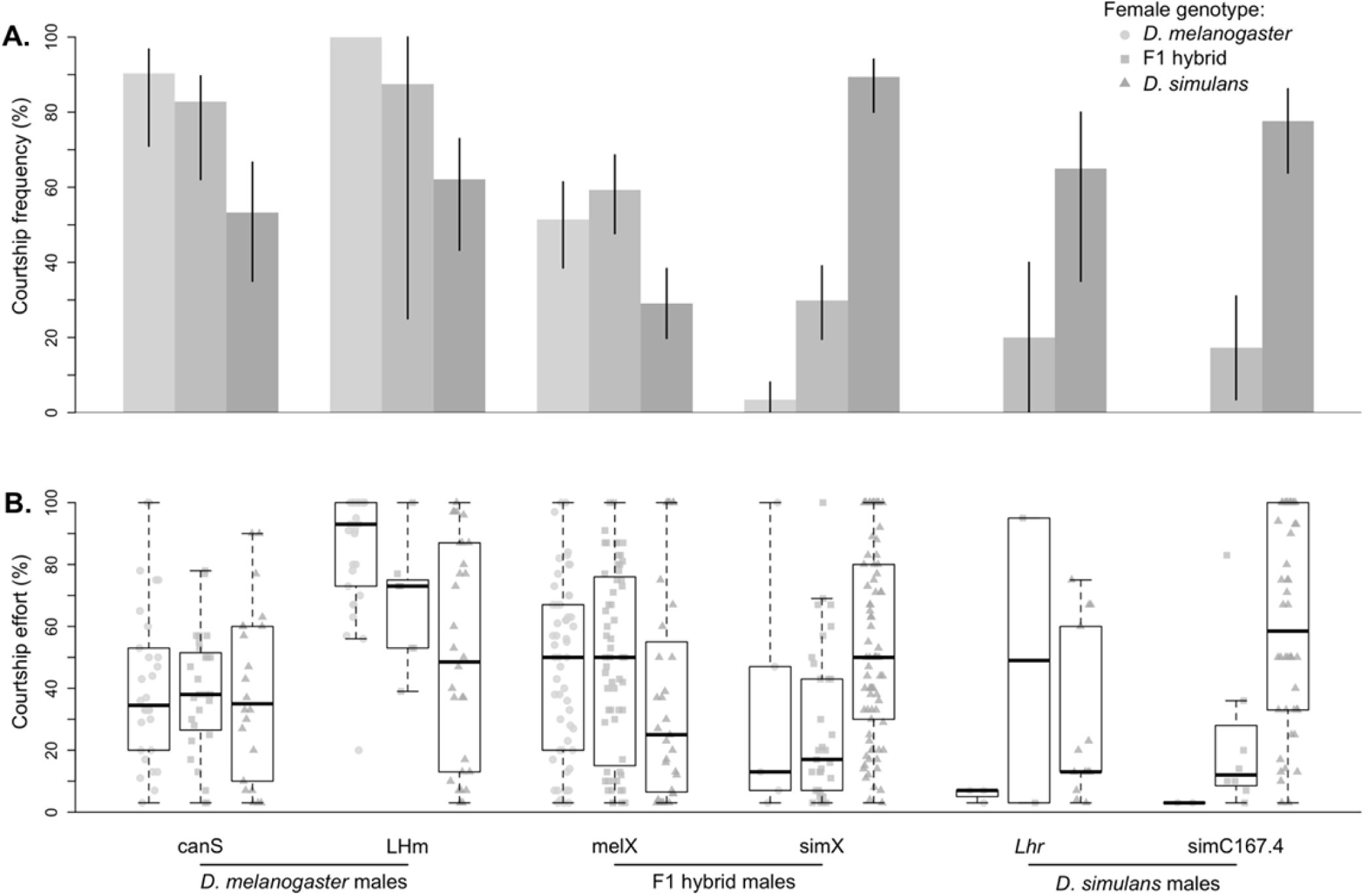
Courtship behaviors of *D. melanogaster* males, *D. simulans* males, and their hybrids. **A.** Courtship frequencies are shown for two strains of *D. melanogaster* (Canton-S and LH_M_), two strains of *D. simulans* (*Lhr* and simC167.4), and their reciprocal hybrids (melX and simX) toward three female types: *D. melanogaster* (light grey), *D. simulans* (grey), and F1 hybrids (dark grey). Whiskers represent 95% bias corrected and accelerated bootstrapped confidence intervals. **B.** Courtship effort is shown for the same male and female combinations. Courtship effort is calculated as the percent of time that males spent courting (only males that courted were included in these calculations). Boxplots show the median (bold black line), interquartile range (box) and full extent of the data excluding outliers (whiskers).

We also observed each *D. melanogaster* and *D. simulans* strain with F1 hybrid females (Figure 2A, Table S2). These females still produce 7,11-HD, due to a single functioning copy of *desatF* (Shirangi et al., 2009), and concordantly, *D. melanogaster* and *D. simulans* males court them similarly to *D. melanogaster* females. We found that our *D. melanogaster* strains were just as likely to court F1 hybrid females as they were to court *D. melanogaster* (p = 0.6486 for LH_M_, and p = 6724 for Canton-S), and these comparisons remained non-significant when we calculated a consensus p-value (p = 0.7980). Conversely, for one of our *D. simulans* strains, simC167.4, we found that males court F1 hybrid females significantly less often than *D. simulans* females (p = 5.59E-07). *Lhr* males had a much lower courtship frequency with F1 hybrid females compared to *D. simulans* females (20% vs. 65%, respectively), but this difference was not significant, likely due to small sample size (p = 0.1003 and N = 10). Supporting this, a consensus p-value for both *D. simulans* strains found that *D. simulans* overall had a significantly higher courtship frequency with *D. simulans* females than with F1 females (p = 9.93E-07). For both *D. simulans* strains, we detected no difference in courtship frequency toward F1 and *D. melanogaster* females, as all had relatively low courtship frequencies (for *Lhr* p = 0.1230, and for simC167.4 p = 0.2089; and consensus p = 0.1197).

We find less striking differences when we calculate the courtship effort of those males that did court (Figure 2B, Table S2). For example, once they began courting, Canton-S males courted all three female types with equal vigor (p = 1 for all comparisons). LH_M_ males, however, courted *D. melanogaster* with much higher vigor than *D. simulans* females (p = 0.0011). Thus, unlike courtship frequency, courtship effort appears to have strain-specific effects within the *D. melanogaster* strains we surveyed. For *D. simulans*, we detected a nearly significant increase in courtship effort for simC167.4 males that courted *D. simulans* females compared to males that courted F1 females (p = 0.0506). We are unable to detect significant differences in courtship effort for *Lhr* males among any comparisons, or for simC4 male comparisons involving *D. melanogaster* females, because so few males courted *D. melanogaster* females (and F1 females for *Lhr*).

### The X-chromosome partially explains differences in courtship behavior

Qualitatively, the behavior of hybrid males largely replicates the behavior of the X chromosome donating parent (Figure 2, Table S2). Like males of the *D. melanogaster* parent strain (LH_M_), melX hybrid males court both *D. melanogaster* and F1 hybrid females significantly more often than *D. simulans* females (*D. melanogaster*: p = 0.0052, F1: p = 0.0005 Figure 2A). Also like the LH_M_ parent strain, melX males court F1 hybrid females and *D. melanogaster* females at similar frequencies (p = 0.6724). Quantitatively, however, the behavior of melX males does not entirely replicate that of the *D. melanogaster* parent strain. For instance, although still significantly higher than when paired with *D. simulans* females, melX males court both *D. melanogaster* females and F1 females at significantly lower frequencies than LH_M_ (p = 9.92E-07 and p = 0.0022, respectively). With respect to courtship effort, melX males court each female indistinguishably (all p > 0.8541, Figure 2B).

The behavior of simX males more closely reproduces that of the *D. simulans* parent strain (*Lhr*). Like *Lhr*, simX males court *D. simulans* females at much higher frequencies than *D. melanogaster* females (p = 1.32E-15, Figure 2A) and F1 females (p = 4.03E-14). Unlike the *Lhr* parent strain, simX males court F1 females significantly more frequently than *D. melanogaster* females (p = 1.57E-05). Courtship towards *D. melanogaster* was still too rare to detect differences in courtship effort compared to *D. simulans* or F1 females (both p = 1, Figure 2B), but simX males courted *D. simulans* with significantly higher effort than F1 females (p = 0.0005, Figure 2B). Quantitatively, simX males behave very similarly to their *D. simulans* parents. simX males court *D. melanogaster* and F1 females with frequencies equivalent to that of *Lhr* males (p = 0.5876 and p = 0.7190, respectively), but they court *D. simulans* females at a higher frequency (p = 0.0373). We suspect this latter result is a byproduct of increased heterozygosity relative to the inbred *Lhr* parent strain.

### A single region of the D. melanogaster X chromosome changes simX hybrid courtship behavior

To test specific regions of the X chromosome for their role in courtship preference differences, we measured the courtship behavior of 16 duplication hybrids (Table 1A, Figure 1C). These duplication hybrids are simX hybrids made heterozygous for one stretch of the *D. melanogaster* X chromosome. We observed 15 of these strains with *D. melanogaster* females, and interestingly, none displayed courtship (N = 6-20 for each, N = 179 total, Table S2). All 16 of the duplication hybrid strains did court both *D. simulans* and F1 females, however. We detected significant variation in the amount of courtship these lines displayed to both female courtship targets (χ^2^= 55.36, df = 15, p-value = 1.55e-06 for *D. simulans* females, N = 709 total; and χ^2^= 66.58, df = 15, p-value = 1.80e-08 for F1 females, N = 759 total).

On average, the duplication hybrid lines had significantly higher courtship frequencies toward *D. simulans* females than toward F1 females (average CF = 35.86% for *D. simulans* females, average CF = 19.30% for F1 females, p = 5.43E-08, N = 16). This pattern is consistent with what we see for simX hybrids without X-linked duplications, which similarly courted *D. simulans* females more often than F1 females. However, courtship frequencies with both females are significantly higher for simX males compared to duplication hybrids (average simX CF with F1 females = 30%, Student’s t = -3.3664, df = 15, p = 0.0042; average simX CF with *D. simulans* females = 89%, Student’s t = -12.891, df = 15, p = 1.614e-09). In total, 15 of the 16 duplication hybrid genotypes courted *D. simulans* with higher frequency than F1 females (Figure 3A, Table S2); three of these lines showed a significant preference for *D. simulans* females after correction for multiple tests (p = 0.0415 for BSC296, p = 0.0061 for BSC277, and p = 0.0437, for BSC200). Only one duplication hybrid strain, BSC100, courted F1 hybrids with higher frequency than *D. simulans* hybrids (p = 0.0136). In general, the duplication hybrid strains courted *D. simulans* females with greater effort than F1 hybrid females (grand median CE = 17% for *D. simulans* females, and the grand median CE = 10% for F1 females, p = 8.78E-05). Again, BSC100 duplication hybrids were the only hybrids to display higher courtship effort toward F1 females (CE = 30%) than toward *D. simulans* females (CE = 13.33%), although this difference was not significant after correcting for multiple comparisons (p = 0.0976, Figure 3B, Table S4).

**Figure 3.**
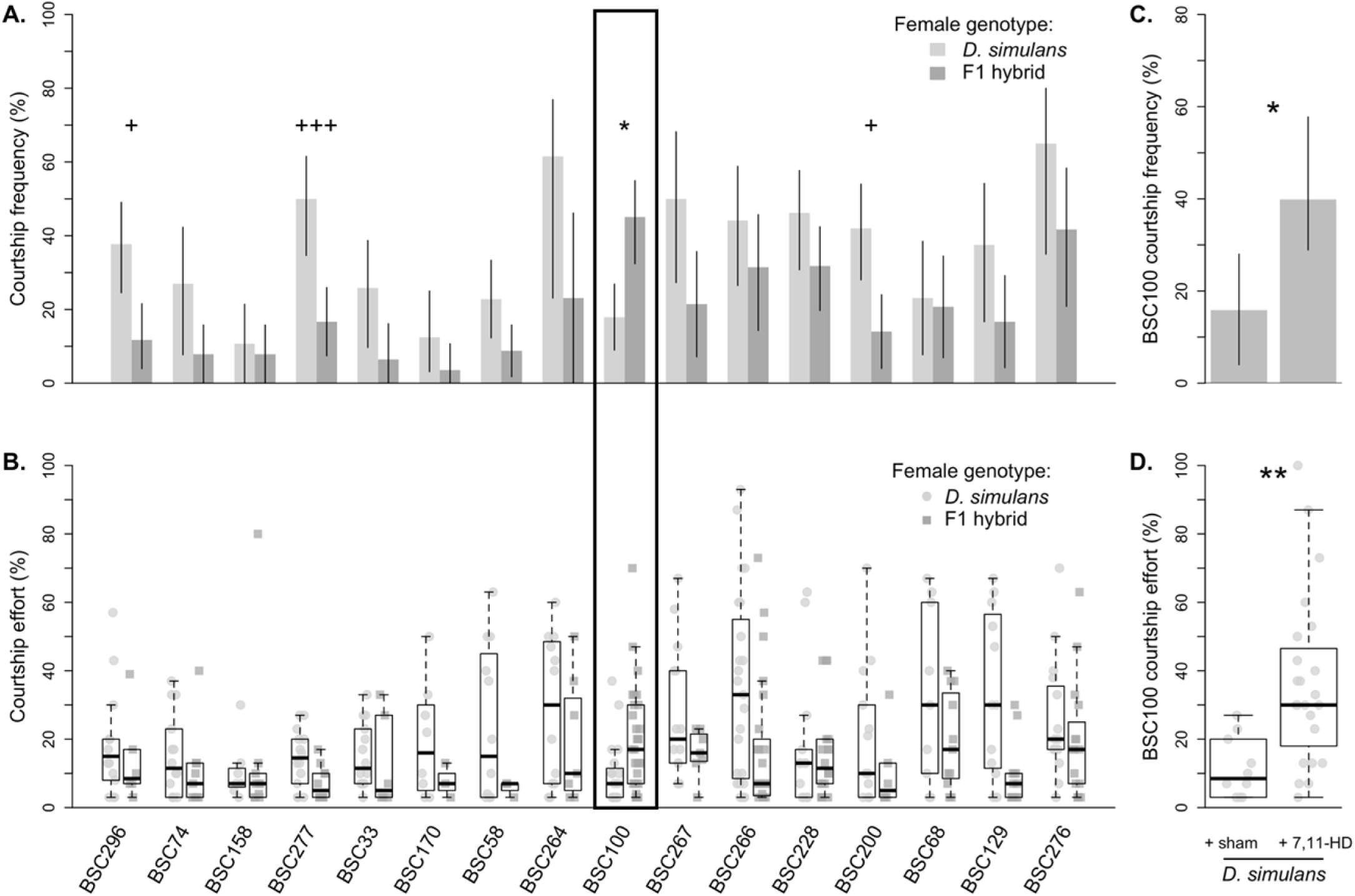
Courtship behaviors of duplication hybrid males. **A.** The courtship frequencies for each of the 16 duplication hybrids when paired with both *D. simulans* females (light grey) and F1 females (dark grey). **B.** The courtship effort for each of the 16 duplication hybrids when paired with *D. simulans* females (light grey circles, left) and F1 females (dark grey squares, right).BSC100, the only duplication to display greater courtship frequency and effort towards F1 hybrids over *D. simulans* females is enclosed within the black box. **C.** The frequency of BSC100 males that courted sham-perfumed *D. simulans* females (left), and *D. simulans* females perfumed with synthetic 7,11-HD (right). **D.** The courtship effort of BSC100 males that courted sham-perfumed *D. simulans* females (light grey circles, left), and *D. simulans* females perfumed with synthetic 7,11-HD (dark grey squares, right). For all, symbols denote degree of significance after correction for multiple comparisons (* = p< 0.05, ** = p < 0.01, and *** = p < 0.001). Asterisks mark duplication hybrid lines that court F1 hybrid females significantly more. Plus signs mark duplication hybrid lines that court *D. simulans* females more. For A. and C., whiskers represent 95% bias corrected and accelerated bootstrapped confidence intervals. For B. and D., boxplots show the median (bold black line), interquartile range (box) and full extent of the data excluding outliers (whiskers).

### BSC100 duplication hybrid males prefer females with the cuticular hydrocarbon 7,***11-HD***

To verify that CHCs are involved in the courtship behaviors that BSC100 duplication hybrid males exhibited, we perfumed *D. simulans* females with 7,11-HD. When we observed BSC100 duplication hybrid males with perfumed *D. simulans* females, we saw a significant increase in courtship frequency relative to sham-perfumed *D. simulans* females (p = 0.0400, Figure 3C, Table S2). BSC100 hybrid courtship frequency increased from 16% with *D. simulans* sham-perfumed females, to 40% with *D. simulans* females perfumed with 7,11-HD. Note, these courtship frequencies are similar to those seen when BSC100 hybrid males were paired with *D. simulans* females (CF = 17.91%, p = 1) and F1 females (CF = 45.07%, p = 0.8603), respectively. Again, the courtship effort data supports this conclusion (Figure 3D). We observed significantly higher courtship effort toward *D. simulans* females perfumed with 7,11-HD (CE = 30%) when compared to sham-perfumed *D. simulans* females (CE = 3%, p = 0.0014). Encouragingly, the courtship effort we observed toward 7,11-HD-perfumed *D. simulans* females is even higher than that of BSC100 hybrid male courtship effort toward F1 females (p = 0.0062). Additionally, BSC100 hybrid males court sham-perfumed *D. simulans* and non-manipulated *D. simulans* with equal effort (p = 0.4451).

### Fine-mapping the BSC100 region of the X chromosome

The BSC100 X chromosome segment spans ∼1.35 Mb, from position 11,557,017 to 12,903,175 of the *D. melanogaster* X chromosome. This locus contains 159 genes (Table S1). To further refine this region, we measured the courtship behavior of 6 additional duplication hybrids heterozygous for various sections of the X chromosome within our region of interest (Figure 4A, Table 1B). The 6 overlapping duplication hybrids had an average courtship frequency of 42% with *D. simulans* and 20.22% with F1 females, comparable to the original 16 duplication hybrids. Five of the strains (BSC47, BSC49, BSC54, BSC315, and BSC317) had higher courtship frequencies with *D. simulans* females than with F1 females (Figure 4B, Table S2), with three (BSC49, BSC315, and BSC317) being significant (p = 0.0002, p = 0.0462, and p = 0.0057 respectively). Overall, the overlapping duplication hybrid strains also displayed significantly higher courtship effort toward *D. simulans* females (CE = 20%) than toward F1 females (CE = 12.5%, p = 0.0029). The same three strains (BSC49, BSC315, and BSC317) that courted *D. simulans* females with higher frequencies than F1 females also displayed higher courtship effort towards *D. simulans* females (p = 0.0460, p = 0.0408, and p = 0.0408, respectively).

**Figure 4.**
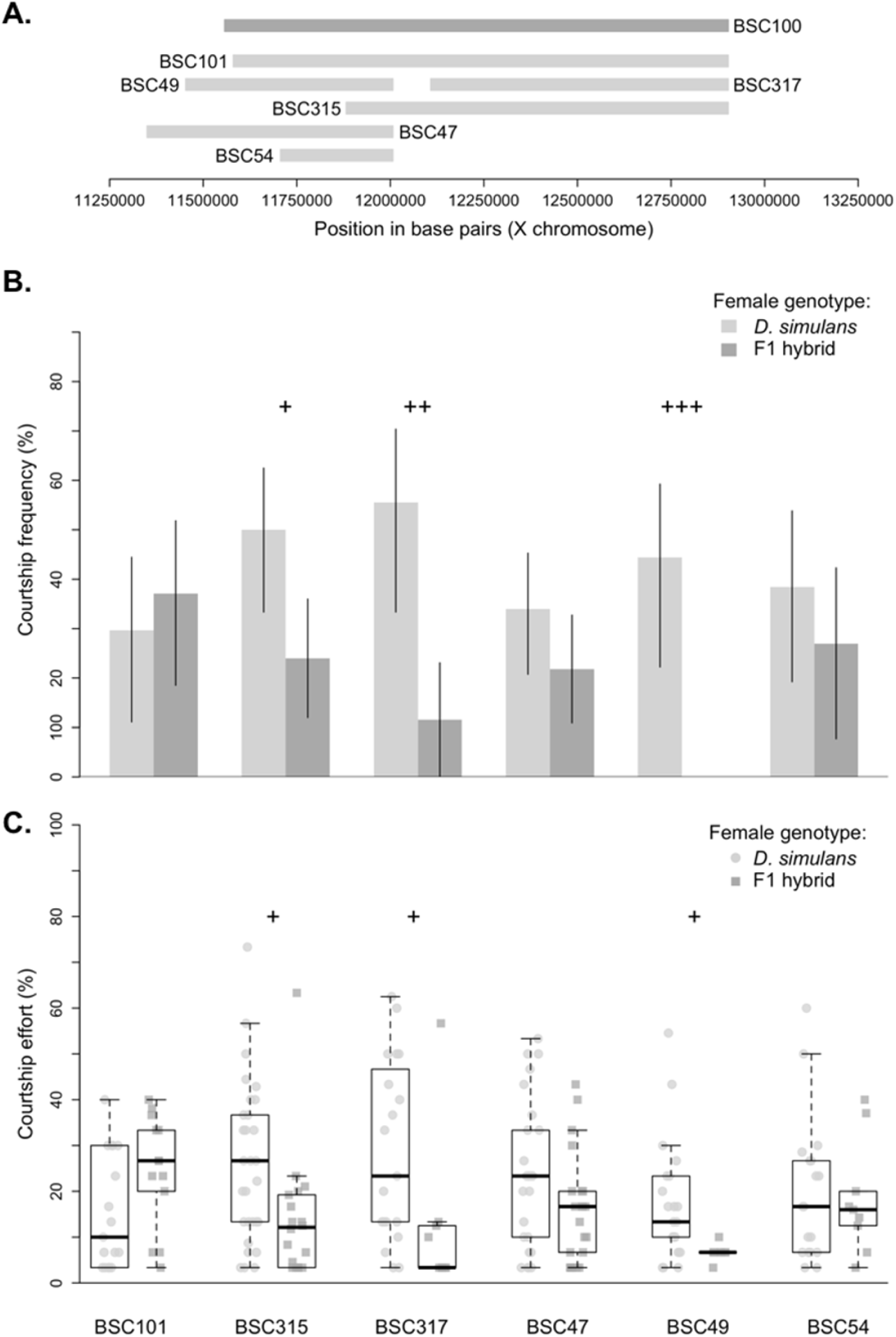
Courtship behaviors of BSC100 overlapping duplication hybrid males. **A.** The physical positions of the original duplication hybrid (BSC100), and the six partially overlapping strains we assayed to fine-map the region. **B.** The courtship frequencies for each of the 6 overlapping duplication hybrids when paired with both *D. simulans* females (light grey) and F1 females (dark grey). Whiskers represent 95% bias corrected and accelerated bootstrapped confidence intervals. **C.** The courtship effort for each of the 6 overlapping duplication hybrids when paired with *D. simulans* females (light grey circles, left) and F1 females (dark grey squares, right). Boxplots show the median (bold black line), interquartile range (box) and full extent of the data excluding outliers (whiskers). For all, plus signs mark duplication hybrid lines that court *D. simulans* females more, and denote degree of significance after correction for multiple comparisons (+ = p< 0.05, ++ = p < 0.01, and +++ = p < 0.001).

A single strain, BSC101, displayed a different pattern than the others. Interestingly, this duplication segment spans 98% of the region covered by BSC100, while the other duplications cover various smaller portions of the region, ranging from 22% to 76% coverage (Figure 4A). BSC101 had a higher courtship frequency when paired with F1 females (CF = 37.04%) than when paired with *D. simulans* females (CF = 29.63%), though not significantly so (p = 1 after correcting for multiple comparisons). However, BSC101 duplication hybrids also did not differ from BSC100 duplication hybrids with respect to courtship frequency towards either *D. simulans* or F1 females (p = 0.2663 and p = 0.5028, respectively). Likewise, BSC101 duplication hybrids were the only strain to display higher effort when paired with F1 females (CE = 26.67%) than when paired with *D. simulans* females (CE = 10%), though not significantly so (p = 0.2067). These numbers are also comparable to the courtship effort of BSC100 males towards each female type.

### Testing candidate genes

The BSC100 region is known to contain 159 genes (Table S1). To reduce this list to a testable number, we focused on genes with neurological functions and *fruitless* binding sites (see Methods). This produced 6 candidate genes: *papillote* (pot), *cacophony* (cac), *Tenascin-a* (*Ten-a*), *Smrtr* (*Smr*), *Ionotropic receptor 11a* (*Ir11a*), and *Phosphodiesterase 9* (*Pde9*, Figure 5). We used loss-of-function mutations or RNAi to investigate 5 of these 6 genes further (unfortunately, *pot* loss-of-function hybrid males were inviable).

**Figure 5.**
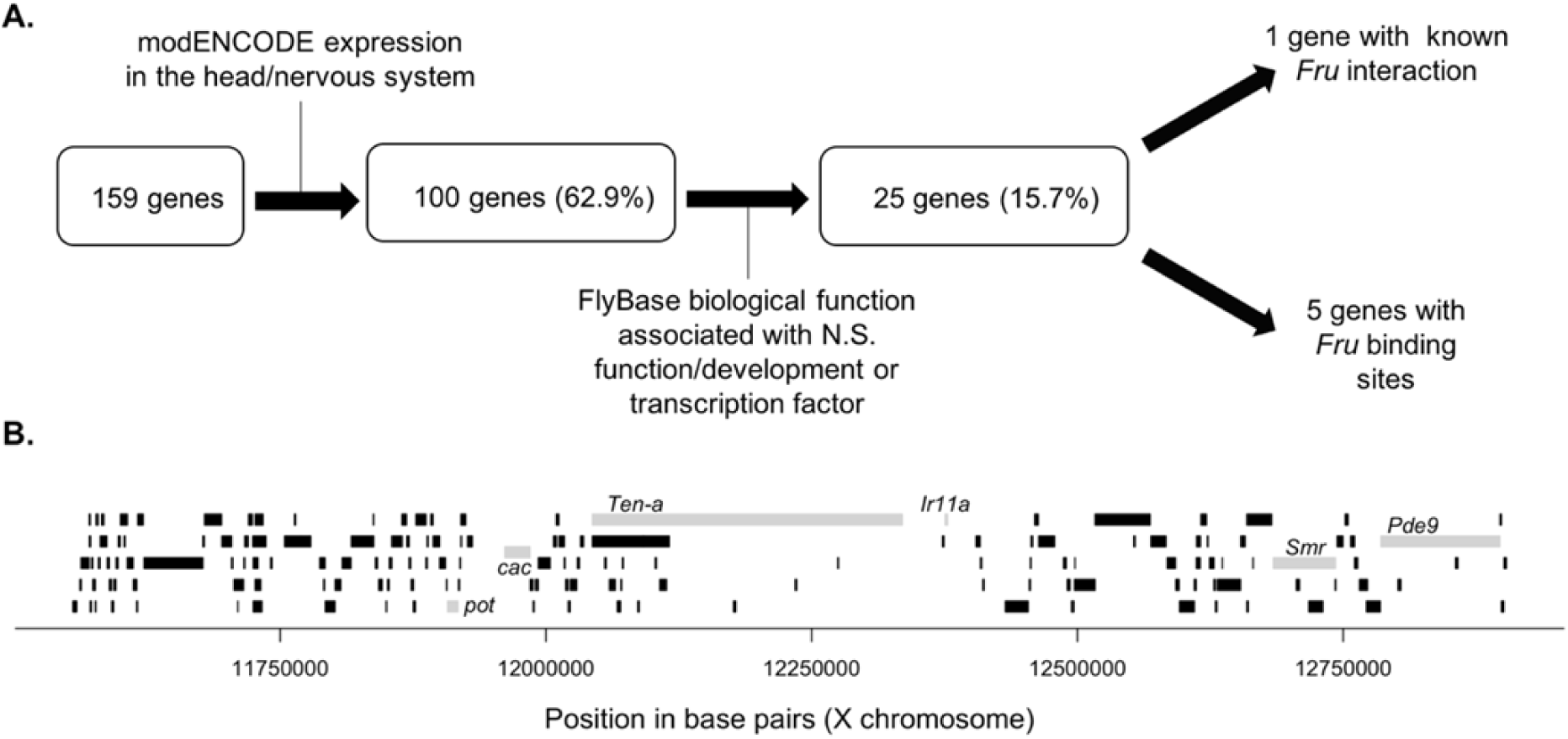
The 159 genes uncovered by BSC100. **A.** A pipeline for identifying relevant candidate genes among the 159 covered by BSC100. We expect genes to be expressed in the central nervous system and to have nervous system functions (or act as transcription factors for loci with such functions). Because *Fruitless* is important to wiring the male nervous system, we also looked for specific genes that interact with *Fru* as particularly strong candidates. **B.** A schematic of the 150 genes. The 6 genes that meet our filtering criteria are relatively evenly distributed across the region and shown in grey.

For *Smr*, the hybrid cross between the *D. melanogaster* knockout strain produced balancer hybrid (melX males with the X chromosome intact) and *Smr*^-^ hybrid males (melX males with the X chromosome lacking a functional *Smr* gene). *Smr* balancer hybrids courted *D. simulans* and *D. melanogaster* at similar frequencies (p = 1, Figure 6A, Table S2), and with similar efforts (p = 0.1136, Figure 6B). The same is true for *Smr*^-^ hybrid males (p = 1 for both CF and CE). However, these males courted both females at significantly higher frequencies than balancer hybrids (p = 0.0073 for *D. simulans*, and p = 0.0135 for *D. melanogaster*). Despite quantitative differences, both *Smr*^-^ and balancer hybrid males show no preference for *D. melanogaster* or *D. simulans* females. Thus, there is no clear effect of the loss of *Smr* on male preference when compared to intact balancer hybrid males.

**Figure 6.**
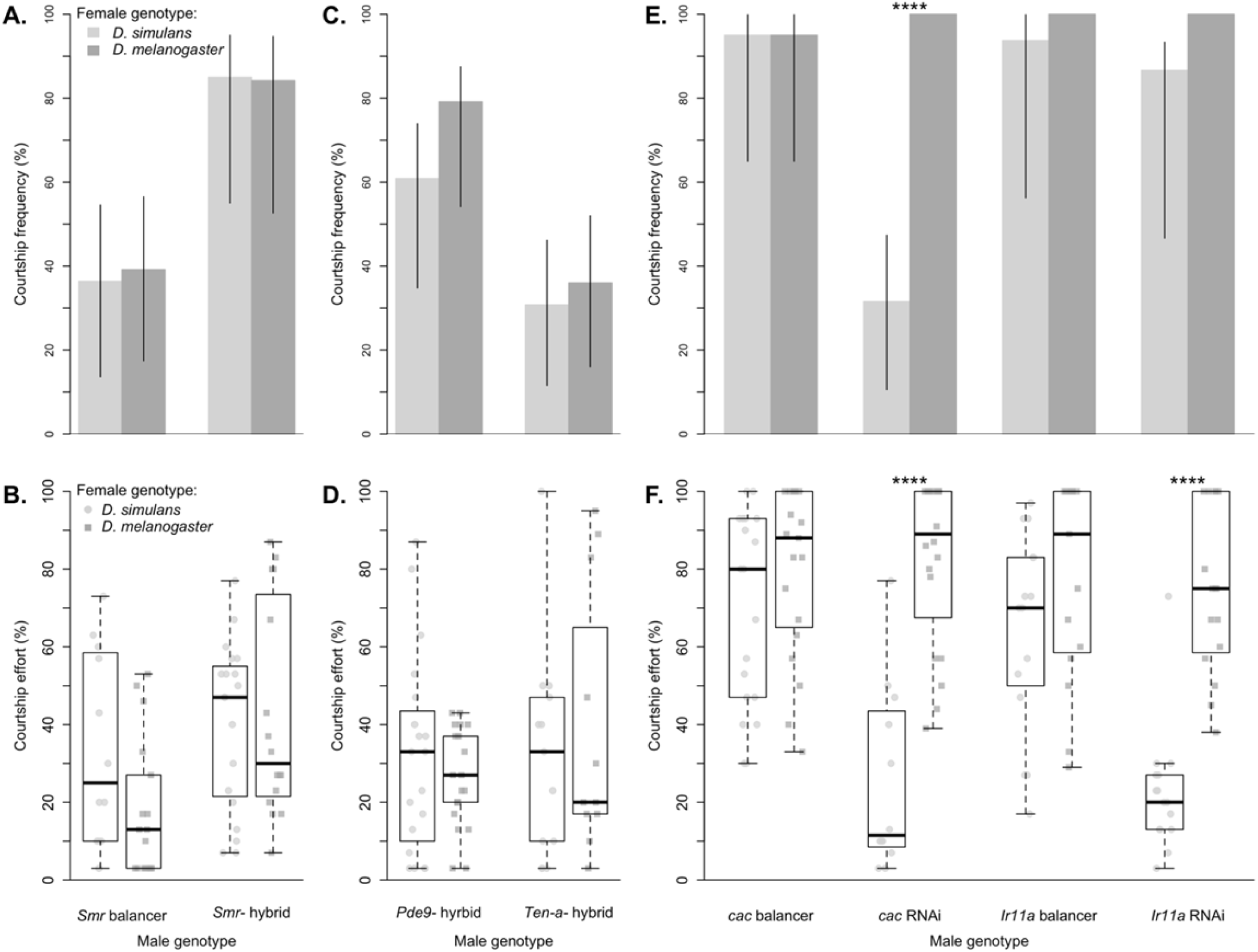
The courtship behaviors of RNAi knockdown and *D. melanogaster* X chromosome hybrids with individual gene knockouts. **A.** The courtship frequency of *Smr* knockout hybrid males (*Smr* hybrid) is compared to *Smr* X chromosome balancer males (*Smr* balancer). **B.** The courtship effort of *Smr* knockout hybrid males (*Smr* hybrid) is compared to *Smr* X chromosome balancer males (*Smr* balancer). **C.** The courtship frequency for *Pde9* and *Ten-a* knockout hybrid males. **D.** The courtship effort for *Pde9* and *Ten-a* knockout hybrid males. **E.** The courtship frequency of RNAi knockdown and balancer genotypes for two genes, *cac* and *Ir11a* for males paired with *D. simulans* females (light grey bars) and *D. melanogaster* females (dark grey bars). **F.** The courtship effort of RNAi knockdown and balancer genotypes for two genes, *cac* and *Ir11a* for males paired with *D. simulans* females (light grey points) and *D. melanogaster* females (dark grey points). For A., C., and E., whiskers represent 95% bias corrected and accelerated bootstrapped confidence intervals. For B., D., and F., boxplots show the median (bold black line), interquartile range (box) and full extent of the data excluding outliers (whiskers). Asterisks denote degree of significance after correction for multiple comparisons (**** = p < 0.0001).

In contrast, *Ten-a* and *Pde9* are not held over a balancer, so hybrid crosses yield only knockout males. *Ten-a*^-^ hybrids court both *D. simulans* and *D. melanogaster* at equal frequencies (p = 1), and with similar effort (p = 1). *Pde9*^-^ hybrids court *D. melanogaster* at non-significantly higher frequencies than *D. simulans* females (p = 0.2124), and court both females with equal effort (p = 1). In contrast, melX males with functioning copies of these genes court *D. melanogaster* females more frequently (and with non-significantly higher courtship effort). Although these knockout hybrids differ in some aspects from melX males, it is difficult to discern whether these differences are due to the different *D. melanogaster* strains with which these hybrids were made (see Discussion).

For *cac* and *Ir11a*, we compared the behavior of *D. melanogaster* RNAi knockdown flies (Figure 6E,F) to their siblings lacking knockdown (balancer). For *Ir11a*, we found that balancer males court *D. simulans* and *D. melanogaster* indiscriminately in terms of courtship frequency (p = 1) and courtship effort (p = 1). We found the same pattern for *cac* balancer males (p = 1 for both CF and CE). While *Ir11a* RNAi males also courted *D. melanogaster* and *D. simulans* females at equal frequencies (p = 0.9032), *cac* RNAi males had significantly lower courtship frequencies with *D. simulans* females than with *D. melanogaster* females (p = 1.34E-05). Interestingly, both *Ir11a* and *cac* RNAi males displayed reduced effort toward *D. simulans* females compared to *D. melanogaster* females (p = 8.13E-06 and p = 1.76E-05, respectively).

To test if the reduction in courtship frequency and effort of *cac* RNAi males with *D. simulans* female is driven by the absence of 7,11-HD on the *D. simulans* cuticle, we also observed *cac* RNAi and balancer males with sham- and 7,11-HD perfumed *D. simulans* (Figure 7A,B). We found that 7,11-HD does indeed cause this effect: balancer hybrids court sham perfumed and 7,11-HD perfumed *D. simulans* with equal frequencies (p = 1), and there was no significant difference in courtship effort (p = 0.5876). In contrast, *cac* RNAi hybrids court 7,11-HD perfumed D. simulans at significantly higher frequency and effort than sham perfumed females (p = 3.91E-06 for CF and p = 0.0027 for CE). Thus, *cac* RNAi hybrids require 7,11-HD to stimulate high amounts of courtship.

**Figure 7.**
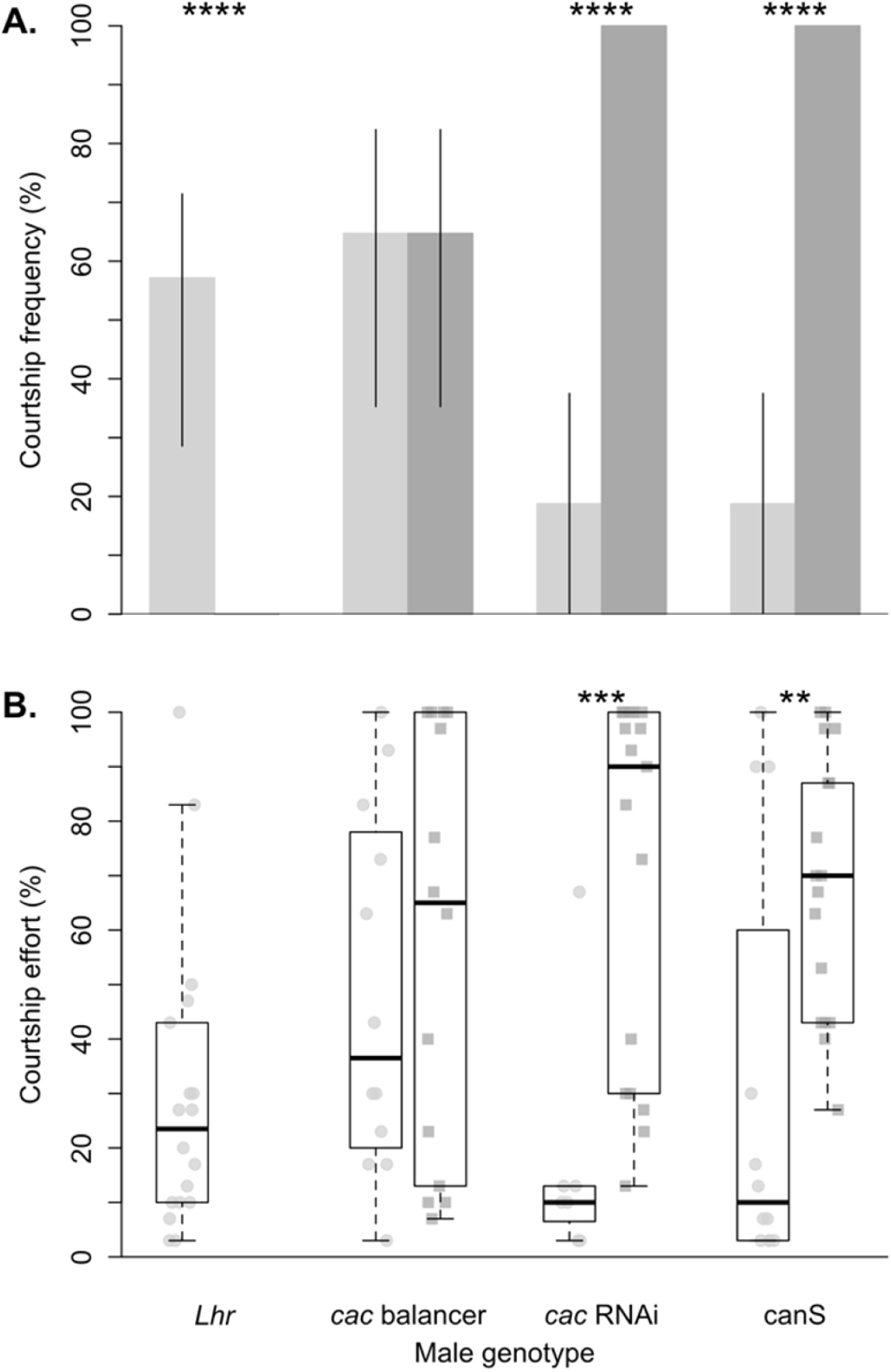
*cac* RNAi male behavior is driven by 7,11-HD. **A.** The courtship frequency of RNAi knock down and balancer genotypes for *cac, Lhr*, and Canton-S (canS) for males paired with *D. simulans* females (light grey bars) and *D. melanogaster* females (dark grey bars). Whiskers represent 95% bias corrected and accelerated bootstrapped confidence intervals. **B.** The courtship effort of RNAi knockdown and balancer genotypes for *cac, Lhr*, and canS for males paired with *D. simulans* females (light grey circles) and *D. melanogaster* females (dark grey squares). Boxplots show the median (bold black line), interquartile range (box) and full extent of the data excluding outliers (whiskers). For all, asterisks denote degree of significance after correction for multiple comparisons (** = p < 0.01, and *** = p < 0.001, **** = p <0.0001).

## Discussion

### An important difference in male courtship preference between D. melanogaster and D. simulans partially maps to the X chromosome

Our observation of two *D. simulans* and two *D. melanogaster* strains confirms a previously described species difference in male courtship preference, where each species dramatically prefers their own females (Manning, 1959). While we did observe some variation in the quantitative amount of courtship among our lines (Figure 2A), the valence of male preference was always consistent within species. Because we also used females from two different lines, this variation in male behavior could be due to individual variation in female CHC quantity (Pardy et al., 2018), variation in other female traits, and/or variation in male preferences (Pischedda et al., 2014). Our results demonstrate that male courtship preferences act as a large reproductive barrier for *D. simulans* and *D. melanogaster*, as has been shown between *D. simulans* and *D. sechellia* (Shahandeh et al., 2018). A detailed understanding of the genetic basis of this preference would prove illuminating, not only with respect to the evolution of behavior, but also with respect to the evolution of reproductively isolating barriers.

The courtship data we collected using reciprocal *D. melanogaster*/*D. simulans* hybrids (melX and simX males) created in a homogenous background and controlled for cytoplasmic inheritance also confirms the significant role of the X chromosome in male courtship preference differences between these species (Kawanishi et al., 1981). Though we didn’t strictly control for an effect of the Y chromosome, it is unlikely to explain our results because hybrids with the *D. simulans* Y behave more like *D. melanogaster*, and hybrids with the *D. melanogaster* Y behave more like *D. simulans.* Because we only used reciprocal hybrids to demonstrate the X-effect, we cannot rule out the potential of transgressive autosomal effects that cannot be detected in a hybrid background (Mittleman et al., 2017). Indeed, although the behavior of our hybrids qualitatively mirrors that of the X-donating parent, quantitative differences in both courtship frequency and effort, particularly between LH_M_ and melX hybrids, suggest an additional role of autosomal loci (likely *D. simulans* dominant, as simX males behave both qualitatively and quantitatively similar to *D. simulans* males). These results mirror findings from a QTL study mapping male courtship frequency differences among *D. simulans* and *D. sechellia*, another species where females express 7,11-HD and males are stimulated to court by it (M.P. Shahandeh & T.L. Turner, in prep). In this case, the *D. simulans* X chromosome contributes to 7,11-HD aversion, and autosomal *D. sechellia* loci contribute to 7,11-HD attraction. However, in *D. simulans-D. sechellia* reciprocal hybrid males, the effects of the autosomal loci are far greater than the X chromosome, as both hybrids behave more similarly to *D. simulans* males. It remains to be seen whether the same loci affect 7,11-HD response between these species, but the effects/interactions of loci are undoubtedly different.

### A single segment of the D. melanogaster X chromosome has a large effect on male pheromone preference

We expect all DP(1;Y) hybrid strains to behave like the simX hybrid strain, unless the duplicated *D. melanogaster* X chromosome segment harbors dominant or additive male preference loci, in which case they should behave more like melX hybrids. When paired with *D. melanogaster* females, however, we observed no courtship from the 15 DP(1;Y) hybrid strains we observed, which all behaved like simX males in this respect (the 16^th^ unobserved duplication hybrid strain’s CF and CE for *D. simulans* and F1 hybrids also follows this trend). This result is consistent with several possibilities, including: (1) the loci for male courtship preference reside in the 20% of the X chromosome not covered by the duplication hybrid strains, (2) *D. melanogaster* courtship preference alleles are recessive to, or epistatic with, *D. simulans* alleles, the presence of Y-linked X translocations disrupts male behavior in general by making regions of the X chromosome heterozygous, and/or (4) the *D. melanogaster* allele is not expressed properly due to the genomic environment of the translocation. This last possibility is perhaps unlikely, as the duplications have been shown to rescue the loss of 94% of X-linked mutations (Cook et al., 2010).

Similar to our duplication hybrids, we never observed any of our *D. simulans* strains courting *D. melanogaster* females, but each courted F1 females at low levels (18% of all *D. simulans* males courted F1 females), suggesting they may display intermediate female cues. However, *D. simulans* males still strongly preferred *D. simulans* females over F1 hybrid females, likely due to the presence of 7,11-HD on the F1 female cuticle (Coyne, 1996), which suppresses courtship in *D. simulans* males (Billeter et al., 2009). This is also likely the reason that *D. melanogaster* males court F1 females comparably to *D. melanogaster* females. As with the male preference difference between *D. melanogaster* and *D. simulans* females, the preference difference between *D. simulans* and F1 hybrid females still maps to the X chromosome: simX males behave like *D. simulans* males – courting F1 females infrequently, and significantly less frequently and with less effort, than *D. simulans* females (Figure 2). Likewise, melX males behave like *D. melanogaster*, courting F1 females with a similar frequency as *D. melanogaster* females, and significantly more frequently than *D. simulans* females. Because the overall preference patterns between *D. melanogaster* and *D. simulans* were replicated when we compared melX and simX male courtship towards F1 and *D. simulans* females, we additionally observed each of the 16 DP(1;Y) hybrid strains with F1 hybrid females. Again, we expect the courtship preferences of all DP(1;Y) hybrid strains to resemble the simX hybrid strain unless the duplicated *D. melanogaster* X chromosome segment harbors male preference loci, in which case we expect them to court F1 hybrid females at higher frequencies than *D. simulans* females, as we see for melX males.

When we observed hybrid males from the 16 DP(1;Y) genotypes, we found that some males from every strain courted F1 females. Most genotypes courted F1s at low levels, as we saw for simX hybrids. BSC100 hybrids were the only duplication hybrids that displayed higher courtship frequency and effort with F1 hybrids than with *D. simulans* females, replicating the pattern seen in melX hybrid males. However, like simX hybrids, BSC100 hybrids showed no courtship towards *D. melanogaster* females. The fact that BSC100 hybrid males prefer F1 females to *D. simulans* females, but are still unwilling to court *D. melanogaster* females, suggests that the *D. melanogaster* variants at this locus are insufficient to completely mask the effects of the *D. simulans* X genome, which is also present in the BSC100 hybrid. We hypothesize that the greater courtship we see towards F1 hybrid females is not seen towards *D. melanogaster* females because the multiple, partially redundant cues that influence male courtship in *Drosophila* are more intermediate in F1 females (Arbuthnott et al., 2017). If there are multiple male preferences and female cues that have evolved, the BSC100 duplication may recover the *D. melanogaster* preference towards one signal, but still be insufficient to activate the P1 courtship command neurons because of inhibition on other sensory channels by the *D. simulans* genome (Clowney et al., 2015).

This hypothesis is supported by our perfuming data. Although past work has shown that the male preference difference between *D. melanogaster* and *D. simulans* is primarily dictated by female pheromones (Manning, 1959) – specifically 7,11-HD (Billeter et al., 2009) – we could not be sure that that BSC100 hybrid males were responding to this cue, especially because they were unwilling to court *D. melanogaster* females that also express this pheromone. However, we found that BSC100 hybrid males significantly preferred *D. simulans* females perfumed with 7,11-HD to sham-perfumed females, and courted them at a frequency and effort comparable to what we saw with F1 hybrid females, which also have 7,11-HD on their cuticle. These results confirm the role of this X region in 7,11-HD response. Taken together, our findings demonstrate that a single 1.35 Mb segment of the X chromosome has a specific effect on the evolved 7,11-HD preference differences between *D. simulans* and *D. melanogaster*.

### Subdividing this region for fine-mapping results in the loss of the significant preference difference

In order to further fine map the X chromosome region duplicated in BSC100 hybrids, we created 6 additional hybrid genotypes with partially overlapping duplicated segments (Table 1B). When we observed these overlapping duplication hybrid strains, none showed the same pattern we observed for BSC100. Five of these had higher courtship frequencies with *D. simulans* females than with F1 females, just like simX males, although two of these differences were not significant. One strain, BSC101, did have a marginally higher CF and CE with F1 females than *D. simulans* females, albeit non-significantly. This segment has the largest overlap, covering 98% of the segment in BSC100 hybrids (Figure 3C). We hypothesize 2 potential explanations for the loss of significant preference when this region was subdivided.

#### (1) The genetic architecture of male courtship preference within this region is polygenic

It is possible that male courtship preference differences are polygenic— even if these genes are constrained within a single 1.35 Mb segment. In this case, subdivision of this locus may reduce the behavioral effect if these loci are additive, or result in its loss altogether if these loci have epistatic interactions. This possibility fits somewhat with the pattern we observe with our overlapping duplications: hybrids with the smaller overlapping segments have lost the phenotype entirely, while the largest overlap appears to, at least partially, reproduce the phenotype. The simplest model that fits this scenario consists of at least two interacting loci, at either end of BSC100, such that all loci are never captured by any of the overlapping duplication hybrid strains. BSC101 overlaps 98% of BSC100. If this is indeed the case, then at least one locus must reside within the 2% not covered by BSC101. This type of genetic architecture is not uncommon. Many loci contributing to a single phenotype constrained within a single region have similarly been discovered for morphological traits in *Drosophila* and other organisms (Fanara et al., 2002; Harbison et al., 2004; Miller et al., 2014; Peichel & Marques, 2017).

#### (2) Hybrid males heterozygous for regions of the X-chromosome behave differently than typical hybrids

Overall, we observed reduced levels of courtship among our duplication hybrid strains compared to simX or melX hybrids, suggesting that the duplication hybrids behave differently from typical hybrids. The consistent reduction in courtship by duplication hybrids also reduces our statistical ability to detect a significant effect. We feel that abnormal behavior of duplication hybrids – particularly those carrying smaller subdivisions of the initial 16 duplication segments, is the most likely explanation for why our attempts to fine-map the BSC100 region were unsuccessful.

Duplication hybrid males may behave differently for a variety of reasons; the most likely are those that stem from the Y-translocated X-duplication segments themselves. Males made heterozygous for regions of the X chromosome that are typically hemizygous may have abnormal behavior due to epistatic interactions between X chromosome loci. In the melX and simX hybrids, *D. melanogaster* and *D. simulans* X loci are not present in the same genetic background, but they are in duplication hybrids. These loci may interact in unpredictable, non-additive ways, having unforeseen effects on behavior. Alternatively, genes translocated from the X to the Y chromosome may have altered expression patterns due to their new genomic environment, producing a similar effect. Smaller segments, like those we used for fine-mapping, may be more susceptible to this problem, as genes contained within smaller translocated segments are more likely to be surrounded by a foreign genomic environment.

The panel of Y-linked X duplications we used to create duplication hybrids was also created using irradiation (Cook et al., 2010). In fact, each breakpoint was induced by irradiating males, originally creating strains that contained large subdivisions of the X chromosome, like BSC100. Further subdivision of these regions (to create strains like BSC101 and all of the smaller subdivisions that we used for fine-mapping) required additional irradiation. This additional irradiation likely introduced new mutations to the genetic background of these flies, making comparisons between BSC100 and its subdividing strains, like BSC101, imperfect.

Finally, these duplication segments are marked with a dominant visible eye mutation, *Bar*, that substantially reduces the shape of the eye to a small sliver in males. These males likely have restricted fields of vision, and may have difficulty tracking females in the courtship arena, as has been shown for mutations affecting eye pigmentation (Connolly & Cook, 1973; Spiess & Schwer, 1978). Indeed, our data collectors noted during courtship observations that males often seemed to lose track of the females they were courting, and courtship would cease. This, too, likely contributed to the reduced courtship frequency and effort we observed, but is constant across all duplication hybrids. Regardless of the cause(s) of atypical behavior in our duplication hybrids, our failure to fine-map the BSC100 region must be considered in light of the above caveats.

### Testing five candidate genes yields inconclusive results

We were able to test five candidate genes we identified within the BSC100 region, either through the use of gene aberrations or RNAi knockdown. Qualitatively, our experiments using an *Smr* gene aberration suggest that the loss of *Smr* expression in a melX hybrid background has no effect on courtship behavior, as *Smr*^-^ hybrids behaved like *Smr* balancer hybrids in that they court both *D. simulans* and *D. melanogaster* females at equal frequency and with equal effort. Quantitatively, however, *Smr*^-^ hybrids showed higher overall courtship frequencies and efforts towards both females compared to balancer hybrids. This result suggests that males harboring an X chromosome balancer are less vigorous, and may behave atypically due to the presences of large inversions on a hemizygous sex chromosome. Thus, balancer hybrids are not an ideal comparison.

The results of our comparisons of *Ten-a*^-^ and *Pde9*^-^ hybrids are congruent with that of *Smr*^-^ hybrids. *Ten-a*^-^ and *Pde9*^-^ hybrids also court both female types with equal frequency and effort. There are, however, quantitative differences in the courtship frequencies of each hybrid, suggesting that *D. melanogaster* genetic background also influences male behavior (each has the same *D. simulans* background), making comparisons between strains imperfect.

Nonetheless, in courting indiscriminately, all three of these strains display a different overall pattern of courtship than intact melX males, which court *D. melanogaster* females at significantly higher frequencies than *D. simulans* females (melX males also display non-significantly higher efforts with *D. melanogaster* females). This may simply be because melX males were created using a different *D. melanogaster* background (LH_M_), and other *D. melanogaster* backgrounds may not discriminate as strongly (consistent with the *Smr* balancer hybrid data and RNAi/balancer data discussed below). In the case of *Ten-a*^*-*^ hybrids, the *D. melanogaster* background is Canton-S, which also courts *D. melanogaster* more frequently than *simulans* females, and does not differ from LH_M_ in this respect (Figure 2A). Thus, it may also be because each of these genes plays a small roll in reducing male preference for *D. melanogaster* females, and additively produce a much larger effect, like that seen with BSC100 hybrids.

The results of our RNAi knockdown screen did identify one gene that, when knocked down, significantly changed male behavior: *cac*. Although there was a general effect of RNAi knockdown on courtship effort overall, only *cac* RNAi males showed reduced courtship frequency towards *D. simulans* females, while *Ir11a* RNAi males and both *cac* and *Ir11a* balancer males displayed high courtship frequencies to both species. Further, we’ve shown that this effect is driven by the lack of 7,11-HD on the *D. simulans* female cuticle, because when we add 7,11-HD synthetically, *cac* RNAi male courtship frequency and effort with *D. simulans* females return to high levels. This result is, at first, not entirely intuitive; the loss of expression of a *D. melanogaster* allele makes males behave comparatively *more* like *D. melanogaster*. We take this result to imply that *cac*, a calcium channel subunit expressed in neurons, is important to general signal transduction in the male CNS. Thus, the loss of *cac* expression results in difficulty activating P1 courtship neurons in the absence of multiple attractive stimuli, like 7,11-HD, that activate different pathways converging on P1 courtship neurons (Clowney et al., 2015). If anything, this result suggests that *cac* is neither necessary nor sufficient for 7,11-HD response.

Ultimately, these results are difficult to interpret, because it is unclear what phenotypes to expect from a hemizygous male harboring a gene disruption. The only case where this test should yield a clear result is where the difference in behavior is attributable to a loss of function or expression in *D. simulans*. However, differences in phenotype can also be due to coding differences among genes, or differences in the amount, location, or timing of gene expression. In these cases, it is unclear what phenotypic change to expect from males completely lacking a gene altogether. While comparing gene aberrations from a *D. simulans* male may provide the reciprocal test (if phenotypic change is due to a loss in *D. melanogaster*), this test is still subject to all of the same problems discussed above. Additionally, it would require significant time and effort to create these aberrations, as none are currently available in *D. simulans* strains. This difficulty is specific to mapping male phenotypes on the X chromosome, as quantitative complementation (Turner, 2014) and reciprocal hemizygosity tests are not possible (Stern, 2014).

It is important to note that we only tested five candidate genes. We selected these genes because they met specific criteria (see methods) that we believed made them likely candidates, and because screening 159 knockouts is too large an undertaking (we observed nearly 500 pairs of courting flies to test our 5 candidates). It is quite possible, then, that the gene(s) responsible are among those that did not meet our strict criteria. For instance, perhaps the gene(s) are expressed in the developing larvae, rather than the adult CNS. Alternatively, perhaps the gene(s) are not specific to the nervous system, and instead have a more general function. Finally, the gene(s) may not directly interact with *Fru*, and instead act downstream or independently of *Fru*.

## Conclusions

Our results demonstrate that male courtship preference differences between *D. melanogaster* and *D. simulans* is at least partially explained by 1.35 Mb region of the X chromosome. We further show that this region responds to the presence of the *D. melanogaster* cuticular hydrocarbon pheromone, 7,11-HD. Unfortunately, attempts to fine-map this region were unsuccessful using the DP(1;Y) hybrid method and a candidate gene approach. Because hybrid offspring of these species are sterile, we cannot pursue other avenues to map the causal loci, such as QTL mapping. Similarly, because males are hemizygous for the X chromosome, we cannot use large engineered chromosomal deletions (as in Laturney & Moehring, 2012; Pardy et al., 2018).

Still, our findings contribute to our understanding of the 7,11-HD preference phenotype. Although the neuronal circuitry required for 7,11-HD response in *D. melanogaster* has been known for some time (Clowney et al., 2015), it was just recently found that the same circuitry detects and responds to 7,11-HD in *D. simulans* (Seeholzer et al., 2018). While the anatomy of this circuit has remained constant during the divergence of these species, the valence of male response has undoubtedly changed – in large part due to changes in the interactions between these neurons, rather than their physical connections. It is still unclear what genetic changes are required to modify the interactions of these neurons, but our results provide a narrowed region of the genome with which to identify and continue to test candidates. Our results also highlight the difficulty of dissecting such a complex phenotype using a purely mapping approach. It is our hope that these data, when paired with functional dissection of the nervous system, can contribute to the identification of alleles explaining behavioral evolution. This is a necessary goal if we wish to understand the common patterns of genetic change underlying behavioral divergence.

## Acknowledgements

We are grateful to Veronica Cochrane, Cameryn Brock, Susanne Tilk, Wesley Cochrane, Maddie Evancie, Devon Cooper, and Eamon Winden for their hours spent observing Drosophila courtship. We also thank the undergraduate students of the MCDB161L M’13 course at UCSB. This program was supported in part by a grant to the University of California, Santa Barbara from the Howard Hughes Medical Institute through the Science Education Program. This work would not have been possible without the community-supported resources available at the Cornell *Drosophila* Stock Center (CSBR 1820594), the Bloomington *Drosophila* stock center (NIH P400D018537), or FlyBase (flybase.org; Gramates et al., 2017). This work was funded by the National Institutes of Health (R01 GM098614).

## Supplementary information

**Table S3.** Crosses of *pot* aberrant *D. melanogaster* to *D. simulans* yields no offspring. We crossed *D. melanogaster pot* aberrant females to *D. simulans Lhr* males using the crossing methods described in the main text three separate times. This table shows the offspring count for female, balancer male, and pot^-^ male offspring. No cross produced pot^-^ offspring and only two *pot* balancer males resulted from all three crosses.

| **Table S3** | **Offspring #** |  |  |
| --- | --- | --- | --- |
| **Cross**<br>(female x male) | **females** | ***pot*<sup>-</sup> males** | ***pot* bal. males** |
| *pot* x *Lhr* -1 | 43 | 0 | 1 |
| *pot* x *Lhr* -2 | 49 | 0 | 1 |
| *pot* x *Lhr* -3 | 26 | 0 | 0 |

